# Biotechnological exploitation of *Saccharomyces jurei* and its hybrids in craft beer fermentation uncovers new aroma combinations

**DOI:** 10.1101/2021.01.08.425916

**Authors:** Konstantina Giannakou, Federico Visinoni, Penghan Zhang, Nishan Nathoo, Paul Jones, Mark Cotterrell, Urska Vrhovsek, Daniela Delneri

## Abstract

Hybridization is an important evolutionary mechanism to bring about novel phenotypes and may produce new hybrids with advantageous combinations of traits of industrial importance. Within the *Saccharomyces* group, *Saccharomyces jurei* is a newly discovered species and its biotechnological potential has not yet been fully explored. This yeast was found to be able to grow well in synthetic wort and at low temperatures, qualities necessary in good candidates for fermentated bevarages. Here, we analysed its fermentation and aroma profile and created novel non-GMO hybrids between *S. jurei* and *S. cerevisiae* ale yeasts to develop new starter strains with interesting flavours for the craft beer industry. Pilot beer fermentations with specific hybrids showed a good fermentation performance, similar to the ale parent strain, while presenting a better sugar attenuation and a more complex flavour profile. This study exploits the genetic diversity of yeasts and show how inter-specific hybridisation and clone selection can be effectively used in brewing to create new products and to eliminate or increase specific traits.

## 1. Introduction

The brewer yeast is often synonymous with *S. cerevisiae*, a species so ubiquitous and predominant in beer and wine fermentations that its domestication predates microbe’s discovery (Gallone et al., 2016). In fact, *S. cerevisiae* offers outstanding fermentation capability together with the ability to consume the broad range of sugars present in wort and must (Mortimer et al, 2000).

Moreover, throughout their domestication, the *S. cerevisiae* strains employed in beer and wine diverged from the wild strains not only for their performances but also for their aroma profile. Wild strain often possesses traits with impart unusual, often unwanted, notes to the aroma, such as the production of the so-called phenolic off flavours (Gallone et al., 2016).

Nevertheless, the role of the *Saccharomyces* species in the production of fermented beverages is not limited to that of *S. cerevisiae* alone. In particular, natural hybrids between *S. cerevisiae* and at least three other species, namely *S. eubayanus*, *S. uvarum*, and *S. kudriavzevii*, were also isolated from brewing and wine making processes (Alsammar and Delneri, 2020) (García-Ríos et al., 2019) (Krogerus et al., 2018). Hybridization helped greatly to bridge the gap between domestic and wild species while combining to the *S. cerevisiae* backbone traits of industrial relevance, such as cold tolerance and flocculation (Giannakou et al., 2020).

A clear example of the potential of interspecific hybridization is represented by *S. pastorianus,* a *S. cerevisiae* and *S. eubayanus* hybrid, employed in the production of lager beers (Mertens et al., 2015) (Monerawela and Bond, 2018). Albeit less common, *S. cerevisiae* hybrids with *S. uvarum* or *S. kudriavzevii* have been associated with wine and cider fermentations while *S. cerevisiae* × *S. kudriavzevii* and *S. cerevisiae* × *S. uvarum* interspecific hybrids have also been isolated from brewing and winemaking fermentative environments (Krogerus et al., 2018) (García-Ríos et al., 2019). Much work has gone in the generation of novel hybrids, both to improve the fitness and to diversify the aroma profile of industrial strains to accommodate the consumer desire for a more complex taste (Bellon, Schmid, Capone, Dunn, & Chambers, 2013; Krogerus et al 2017).

Recently, a new species in the genus, *S. jurei,* was discovered from *Quercus robur* bark in France. Phenotypic assays revealed *S. jurei* to possess interesting traits for industrial application. In fact, the strains characterized resulted resistant to a variety of stressors, from osmotolerance to high sugar concentrations, while presenting a relative high fitness at low temperature (Naseeb et al., 2017; 2018). Recently, *S. jurei’s* application and performance in baking was assessed together with other cryotolerant species of the *Saccharomyces* genus, opening new avenues in the food market (Magalhães et al., 2021).

The craft beer market is constantly growing and in demand of new beer styles and flavours. More complex and stronger flavours to create beer products with natural unconventional ingredients is highly a consumers’ demand (Jaeger et al., 2020). This work is inspired from the need of expanding our biotechnological applications in the craft brewing industry. Beer yeast strains can bring great complexity and novelties in future beer making and the need of new brewing candidates with unique characters is important. Here we evaluated *S. jurei* as a candidate for industrial applications studying its performance in wort media and providing a comprehensive characterization of its aroma profile through GC-GC-MS analysis. Moreover, successfully constructed non-GMO hybrids between *S. jurei* and a *S. cerevisiae* ale strain, containing different combination of allelic traits. The novel hybrids were used in pilot beer fermentation revealing efficient fermentations, optimal sugar attenuation and different spectrum of flavours. This study shows the efficacy of inter-specific hybridisation to create alternative starter cultures for beer fermentations.

## 2. Materials and Methods

### 2.1. Yeast strains

The strains used in this study are the *S. cerevisiae* ale strains OYL200 (Tropical IPA; Omega Yeast lab; Sc-200) and OYL500 (Saisonstein’s monster, Omega Yeast Lab, Sc-500), *S. jurei* D5088 (NCYC 3947; Sj-88) and D5095 (NCYC 3962; Sj-95)(Naseeb et al., 2017), the *S. cerevisiae* lab strain NCYC505 (Sc-505) (Martini and Kurtzman, 1985) and the *S. cerevisiae* wild isolate 96.2 (Sc-96.2) (Paget et al., 2014).

### 2.2. Hybridisation and hybrids confirmation

*S. jurei* D5095 was crossed with *S. cerevisiae* ale strain OYL200. The tetrads were formed by growing the parental strains in pre-sporulation media (yeast extract 0.8%, bacto-peptone 0.3%, glucose 10%) at 30 °C for 16 h before plating on minimal sporulation medium (1% Potassium acetate, 0.125 % yeast extract, 0.1% glucose and 2% bacto-agar). The sporulation plates were incubated at 20 °C for 7–10 days for the formation of tetrads. Tetrad dissection and spore to spore mating was performed in fresh YPD plates (1% yeast extract, 2% peptone, 2% agar, 2% glucose) using a micromanipulator (Singer Instruments 400MSM). The plates were incubated at 30 °C till colony formation. Then, the colonies were spread plated in fresh YPM plates (1% yeast extract, 2% peptone, 2% agar, 2% maltose) and a single colony was used for DNA extraction. DNA was extracted from an overnight grown culture of yeast strains by using the standard phenol/chloroform method described previously (Fujita and Hashimoto, 2000) with modifications described in (Naseeb et al., 2018). The hybrid status of potential hybrids was confirmed by PCR amplification using species-specific primers and genomic DNA template. Two primer pairs were used for the species-specific multiplex PCR, each targeting a specific part of one of the parental species’ genome. Primers Scer_F (5’-GGTTTTATCTGGCACTCAGGT −3’) and Scer_R (5’-GTTGCTGTTGCTGCAAAGGT −3’) amplify a 417 bp amplicon of the *S. cerevisiae* genome. Primers Sjur_F (5’-CTCAAATGGGAATGCCACCG −3’) and Sjur_R (5’-TCCTGATAGTGGTTGTTGCT −3’) generate a 233 bp *S. jurei*-specific amplicon (see Fig. S3 in the supplemental material). The PCR conditions were as follows: 2 min at 94°C, 35 cycles of 1 min at 94°C, 1 min at 55°C, and 30 s of 72°C, followed by a final cycle of 3 min at 72°C and subsequent cooling to room temperature. Candidate hybrids showing two bands were considered to be interspecific hybrids (see Fig. S3). PCR confirmed hybrids were streaked another six times in YPM plates to ensure strain purity and genome stability. To confirm the diploid nature of the hybrids, the ploidy was estimated by flow cytometry as described in (Haase and Reed, 2002) using Amnis ImageStream X (ISX MKII) multispectral imaging flow cytometer.

### 2.3. Micro-fermentations and culture conditions

Growth characterisation and beer brewing ability for the strains of interest was carried out. Three types of experiments were assessed: Growth was examined in 3 different sugars. YP was prepared with 2% glucose, 2% maltose and 2% maltotriose respectively. In addition, high throughput screening with Dry Malt extract (Browland UK) DME starting gravity (OG) 1037.0 was made by using 104 g/L and boiled for 1 hour. The media was filter to remove any undiluted powder. Also, 10L scale beer fermentation was conducted in home brewing equipment (plastic fermenter, lid and airlock) using Pale wort of OG 1055.4. The equipment and wort for this experiment was provided by Cloudwater Brew Co.

Growth kinetics in the 3 different YP media for all strains were created using BMG LabTech Omega series Microplate Readers. The experiment was held for 72h in 96 well plates with 4 biological replicates and initial OD600 was 0.1 in the well plate volumes of 200μl. Growth was recorded via optical density set at 600nm measuring the absorbance at regular intervals and visualized using R.

The micro-fermentations were carried out using a BioLector® I system (m2p Labs, Germany) with a 48 well FlowerPlate® (MTP-48-B, m2p Labs, Germany). The following settings were programmed: filter; biomass (gain 15), humidity; on (>85% using ddH2O), temperature; 20°C, oxygen supply; 20.85% (atmospheric air), agitation speed; 800 rpm. The total volume of each well was 1500 μL, initial OD600 was 0.1. Scatter light at 620nm was measured every 7.29 min and logged by the BioLector®.The final samples (upon completion of the experiment duration) was analysed by HPLC.

The R package grofit was used to analyse the growth curve data of plate reader and micro-fermentation experiments (Kahm et al., 2010). Heat maps of the growth parameters were constructed using the R shiny app (https://kobchai-shinyapps01.shinyapps.io/heatmap_construction/) using min-max column scaling.

The 10L scale beer fermentations were carried out for 14 days using pale wort (see Table S3 for recipe). Each day the gravity (OG) was measured to evaluate the fermentation rate using a density meter (DMA 48, Anton Paar). Sample from the final fermentation day (14^th^) was analysed for aroma composition using GC/MS.

The presence of the *STA1* gene was detected via PCR as described in Yamashita et al., 1985.

### 2.4. Phenotypic assay test

To determine sensitivity to different growth temperatures, standard YPD and YP-maltose agar plates were incubated at 22°C, 16°C and 8°C. The range of temperatures selected are according to temperatures used in the brewing industry. Strains were grown overnight in liquid YPD and 5 μl of 10 fold serial dilutions were spotted to plates starting with an OD_600_ 0.4.

### 2.5. HPLC and Headspace – SPME GCxGC – TOF-MS

The substrate consumption and alcohol production were determined by ion-exchange HPLC. Prior to the HPLC analysis, the samples were filtered (pore size 0.45 μm). The injection volume was 10 μL, the eluent was 5 mM H2SO4 and the flow rate was 0.6 mL/min. A Bio-Rad Aminex HPX-87H column (Hercules, CA, United States) was used and kept at 60°C. Quantification was achieved using a RID-detector. Quantification of sugars and ethanol was achieved with usage of a calibration curve by plotting the instrument response of known standard concentrations of the compounds analysed.

Aroma composition of the final beer products was determined by HS-SPME-GCxGC - TOF-MS. All samples were kept frozen prior analysis to minimize changes occurring during storage. GC instrumentation, SPME extraction, GC × GC – TOF – MS protocol and data analysis was performed as described in (Carlin et al., 2016). Aroma profiling was visualized using R shiny app on heatmap construction (https://kobchai-shinyapps01.shinyapps.io/heatmap_construction/). A Gerstel MultiPurpose Sampler autosampler (Gerstel GmbH & Co. KGMülheim an der Ruhr Germany) with an agitator and SPME fibre was used to extract the volatiles from the sample vial headspace. The GC × GC system consisted of an Agilent 7890 A (Agilent Technologies, Santa Clara, CA) equipped with a Pegasus IV time-of-flight mass spectrometer (Leco Corporation, St. Joseph, MI). A VF-Wax column was used as first-dimension (1D) column, and a RTX-200MS-column was used as a second-dimension (2D) column. The GC system was equipped with a secondary column oven and non-moving quadjet dual-stage thermal modulator. The injector/ transfer line was maintained at 250 °C. Oven temperature programme conditions were as follows: initial temperature of 40 °C for 4 min, programmed at 6 °C min^-^1 up to 250 °C, where it remained for 5 min. The secondary oven was kept 5 °C above the primary oven throughout the chromatographic run. The modulator was offset by +15 °C in relation to the secondary oven; the modulation time was 7 s and 1.4 s of hot pulse duration. Helium was used as carrier gas at a constant flow of 1.2 mL min^-^1. The MS parameters included electron ionisation at 70 eV with ion source temperature at 230 °C, detector voltage of 1317 V, mass range of m/z 35–450 and acquisition rate of 200 spectra s^-^1. SPME extraction was carried as follows: 5 mL of beer, sonicated for 2 min to remove the foam, were put into 20 mL glass headspace vials, 1.5 g of NaCl were added, the samples were spiked with 50 μl of alcoholic solution of 2-octanol at 2.13 mg L^-^1 and 50 μl of alcoholic solution of ethyl hexanoate at 1.0 mg L^-^1 as internal standards. Samples were kept at 35 °C for 5 min and then extracted for 30 min at 35 °C. The headspace was sampled using 2-cm DVB/CAR/PDMS 50/30 lm fibre. The volatile and semi-volatile compounds were desorbed in the GC inlet at 250 °C for 4 min in splitless mode and the fibre was reconditioned for 7 min at 270 °C prior to each analysis.

#### 2.5.1. Data processing and peak annotation

GCxGC-MS data acquisition and processing were achieved with LECO ChromaTOF (Version 4.71). The processing consists of peak picking, peak annotation, and statistic confirmation. During the peak picking, signals which were just above the noise were taken into account (baseline offset = 1). Minimal expected peak width (on 2nd dimension) for deconvolution was 0.8 s. A peak was defined when at least 5 ions whose signal to noise ratio is above 100 (Stefanuto et al., 2017), can be grouped. A picked peak was annotated by matching its mass spectrum (MS) to the reference spectrum in the database. In this study, used MS databases were NIST/EPA/NIH 11, Wiley 8 and the FFNSC 2. The MS similarity threshold for the peak annotation was 700. Each sample was analysed with three technical replications. Inter-measurements peak alignment was performed based on the retention times (both 1st and 2nd dimensions) and mass spectrum. A minimal MS similarity of 600 was required. An analyte was further examined, only if it can be detected in all the technical replications. An inter-class comparison was performed between sample class and blank class. Fisher ratio threshold was used to eliminate artifact compounds (sorbent bleeding, column bleeding and other possible interferences). The applied significance level was 0.05. Peak identification was then completed by checking the linear temperature programmed retention index (LTPRI), which is available in the NIST RI database.

### 2.6. Sensory evaluation of beer products

Sensory analysis was carried out at Cloudwater Brew’s laboratory by brewing experts. Finished beers from the 10L scale fermentation were used by a tasting panel of 6 people focused on the differences and/or similarities on their aroma, flavour, taste/mouthfeel and overall impression. The samples were blind-coded by using 3-digit codes and presented to the trained assessors in random order and duplicates.

## 3. Results and Discussion

### 3.1. Constructions of diploid non-GMO hybrids between S. jurei D5095, D5088 and ale strains S. cerevisiae OYL200 and OYL500

To expand the genetic and phenotypic diversity of yeasts used in the craft beer industry, we generated 8 new interspecific yeast hybrids between different *S. cerevisiae* and *S. jurei* strains.

A previous study showed that *S. jurei* is able to consume maltose, the main sugar found in wort, on solid media (Naseeb et al., 2017). We determined the growth kinetics of *S. jurei* and *S. cerevisiae* lab, wild and ale strains in liquid media containing maltose or maltotriose, as sole carbon source. All strains were able to utilise maltose and maltotriose as sole carbon source, although at different rates, with *S. cerevisiae* ale and lab strains performing better than *S. jurei* and *S. cerevisiae* wild isolate (Figure S1). Specifically, *S. jurei* D5095 was crossed with the commercial ale strain OYL200 while *S.jurei* D5088 was crossed with the *S. cerevisiae* strain OYL500 to create non-GMO hybrids in order to combine the fermentative ability of an ale strain and generate novel aroma profiles. The genome of the diploid parents possess a significant level of heterozygosis (Naseeb et al., 2018), therefore the hybrids will inherit different random combination of traits leading to phenotypic variation. Both *S. jurei* and S*. cerevisiae* strains sporulate well with an efficiency of 80% (Naseeb et al. 2017) and 98% (this study), respectively. We created eight hybrids (H1-8), via spore-to-spore mating. The hybrid nature of the spores was validated via species-specific PCR (Figure S2). The newly created hybrids were sterile as expected *(i.e.* no viable spores after meiosis; data not shown). The DNA content of the hybrids used in pilot fermentations was analysed via FACS to confirm their diploid nature (Figure S3).

### 3.2. Physiological characterisation of the hybrids

The growth of the hybrids was tested in liquid medium in presence of different sugars and compared to the parents (Figure 1 and Table S1). For YP-maltose, YP-maltotriose and YPD the growth curves were assessed using OD_600_ in plate readers (Fig. 1A); while for synthetic wort, which is a dark coloured medium, the growth was assessed using scatter light in the BioLector (Fig. 1B).

**Figure 1:**
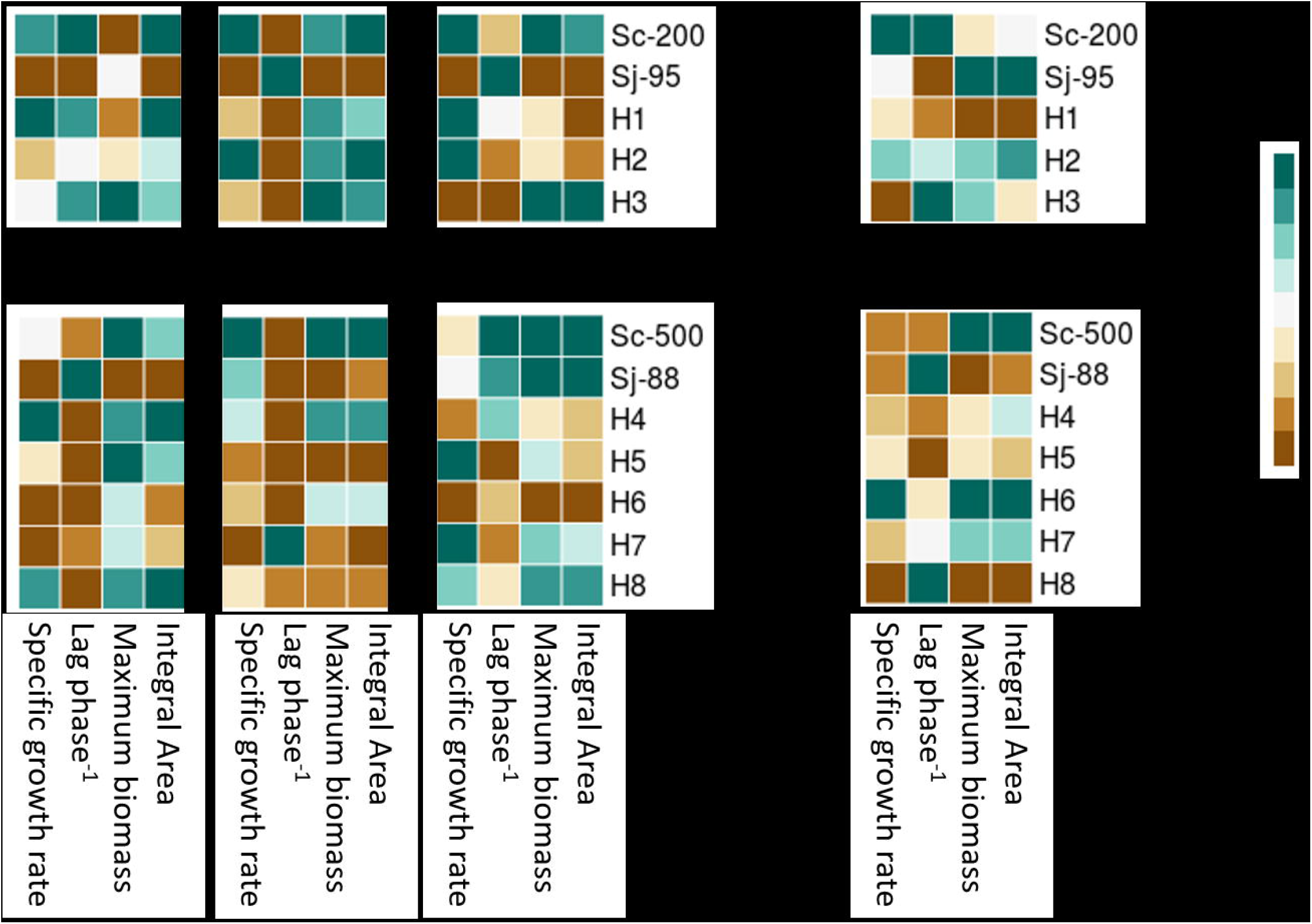
Growth characteristics of *S. jurei* D5095, D5088, *S. cerevisiae* OYL200, OYL500 and H1-8 hybrids. Panel A: 3 different sugar sources at 16 °C after 72h. Panel B: synthetic wort at 20 °C after 72h. For each parameters analysed, max and min represent the best and worst performance respectively for a specific hybrid cross.

In YP + 2% maltose H1 mirror the fermentation performance of Sc-200. H2 and H3 appear to have a slower growth rate and elongated lag phase, but H3 is reaching the highest final biomass among all the strains. In YP + 2% maltotriose a different growing pattern was observed. In this carbon source, H2 is now mirroring the fermentation performance of Sc-200 while, H1 and H3 have moderate performance. H3 continues to have the highest final biomass in this sugar too. In YP + 2% Glucose, all strains behave similarly with small variations in their growth rates. H3 generates the highest biomass from all strains in all 3 sugar sources (Fig.1A).

In conclusion, H1 and H2 performed as well the OYL200 ale parent in maltose and maltotriose, respectively, while H3 reach the highest biomass compared to the other strains in both sugars. Thus, H1 and H2 have inherited different characteristics from the parental ale strain in relation to the efficient utilisation of the different brewing sugars.

Regarding the family of strains Sj-88 × Sc-500 different patterns are observed. In YP + 2% maltose, hybrids H4 and H8 have the maximum growth rate and performing better than the ale parent Sc-500. In YP + 2% maltotriose, however, all hybrids have medium/ low growth rate indicating moderate maltotriose assimilation. H4 is the best among them, in that sugar source, although significantly lower performance than both the parental strains. In YPD, hybrids H5, H7 and H8 had the highest growth rate and greater then ale strain Sc-500.

Overall, from this family, H4 is the best performing hybrid in maltose and maltotriose whilst, H6 is the least good.

Similar results, regarding the performance of the hybrid strains compared with their parents, were observed in the micro-fermentation experiment (BioLector) using synthetic wort, a dry malt extract medium with a high sugar concentration that gives an osmotic stress to the strains (Fig.1B). All the strains derived from the Sj-95 × Sc-200 cross, are able to grow and could cope with the changing environment of sugars, pH and ethanol production. H2 has the faster growth among the hybrids.

For the cross Sj-88 × Sc 500, the hybrid progeny H4, H5 and H6 show higher growth rate than the parent strains. H8 is the worst performing strain in that complex media.

Next, we examined the fitness of all strains in solid media contaning glucose or maltose at different temperatures. As *S. jurei* is coldtolerant (Naseeb et al., 2017), we tested the phenoypes of the hybrids at at 22°C, 16 °C and 8 °C.

Strains from crossings between Sj-95 × Sc-200 in YPD at 22 °C and 16 °C show a similar growth pattern, at 8 °C Sj-95, H1 and H2 grew better compared to the other strains, indicating that traits linked to a higher fitness at cold have been inherited from *S. jurei* (Fig. 2A). In YP-maltose (Fig. 2B) at 8 °C, Sj-95 and H1 grew much better than the other strains. In this medium at 22 °C and 16 °C all the strains grew marginally better than Sj-95.

**Figure 2:**
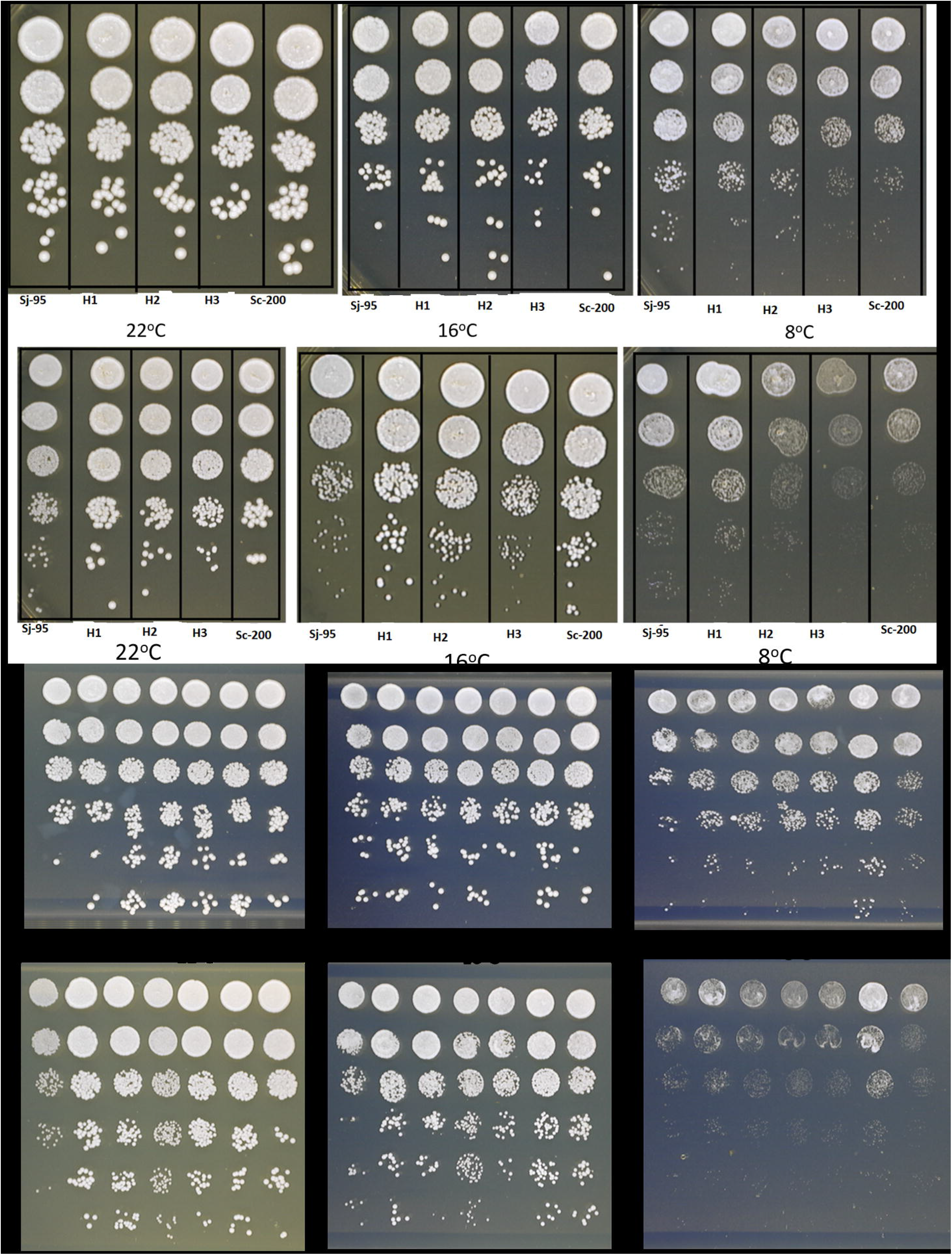
Phenotypic assessment assay plates. Panels A and C: YPD 2% in 22 °C, 16 °C and 8 °C. Panels B and D: YPMaltose 2% in 22 °C, 16 °C and 8 °C. Strains were plated in columns in 10 fold dilutions.

As for the Sc-500 × Sj-88 hybrids, H4-H8, we observe good growth for all strains in YPD at 22 °C and 16 °C. However, at 8 °C H8 has the best growth among all strains (Fig. 2C).

Similarly in YP-Maltose, H8 has the best growth and colony formation among all strains in 8 °C. Whilst, at 22 °C and 16 °C all strains grow well with some small proliferation from H8 and H4 that mirror the ale parent Sc-500 (Fig. 2D).

H1 (crossing Sj-95 × Sc-200) and H8 (crossing Sj-88 × Sc-500) display a good maltose assimilation, inherited from the *S. cerevisiae* ale strain, and a good performance in colder temperatures too as derived from *S. jurei* (Fig. 2).

Ale beer fermentation is taking place between 16-22 °C and overall most of the hybrids grew well in this temperature range, with H1, H2, H4 and H8 showing a better grow in maltose at 16C.

Beer fermentation experiments were then carried out with the hybrid progenies of the cross Sj-95 × Sc-200 (H1-3) based on the performance in DME of the specific Sj parent (Sj −95 is growing better than Sj-88 as shown in Figure S1) and the relative hybrids.

### 3.3. Pilot beer fermentation in 10L vessels

In beer fermentation, sugar attenuation and cell viability are important at the end of fermentation (Sanchez et al., 2012). In brewing terms, attenuation describes the level of wort carbohydrates that are converted in ethanol during fermentation (Vidgren et al., 2009) and it is desired to a level of 70-80% as the leftover sugars do contribute to the final beer products conferring a desirable sweetness and body composition. Certain *S. cerevisiae* strains are characterized by their ability to secrete glucoamylase *(STA1* gene), an enzyme that catalyses the digestion of dextrins. This amylolytic activity can lead to hyper-attenuation, meaning that almost all the sugars (fermentable and non-fermentable) are consumed during beer fermentation, and/or secondary fermentation which can cause excess carbon dioxide formation in bottles, cans or kegs (Meier-Dörnberg et al., 2018) (Yamashita et al., 1985). Results from 10L beer fermentation experiment evaluated the sugar attenuation and cell viability of all strains at pilot scale (Fig. 3). H3 and Sc-200 are over attenuating the beer, while H1 and H2 resulted in approximately 70% attenuation in final beer, a characteristic that is desirable for different beer styles. The gravity remained stable after the 14^th^ day indicating that no further fermentation took place. All the strains were stained with methylene blue to estimate cell viability which is important for re-pitching. No significant difference in viability across strains was detected *(i.e.* approximately 50% viable cells at the end of the fermentation process). *S. jurei* D5095 beer fermentation was not completed as the gravity remained relatively high which is expected from the lab scale growth kinetics we previously observed.

**Figure 3:**
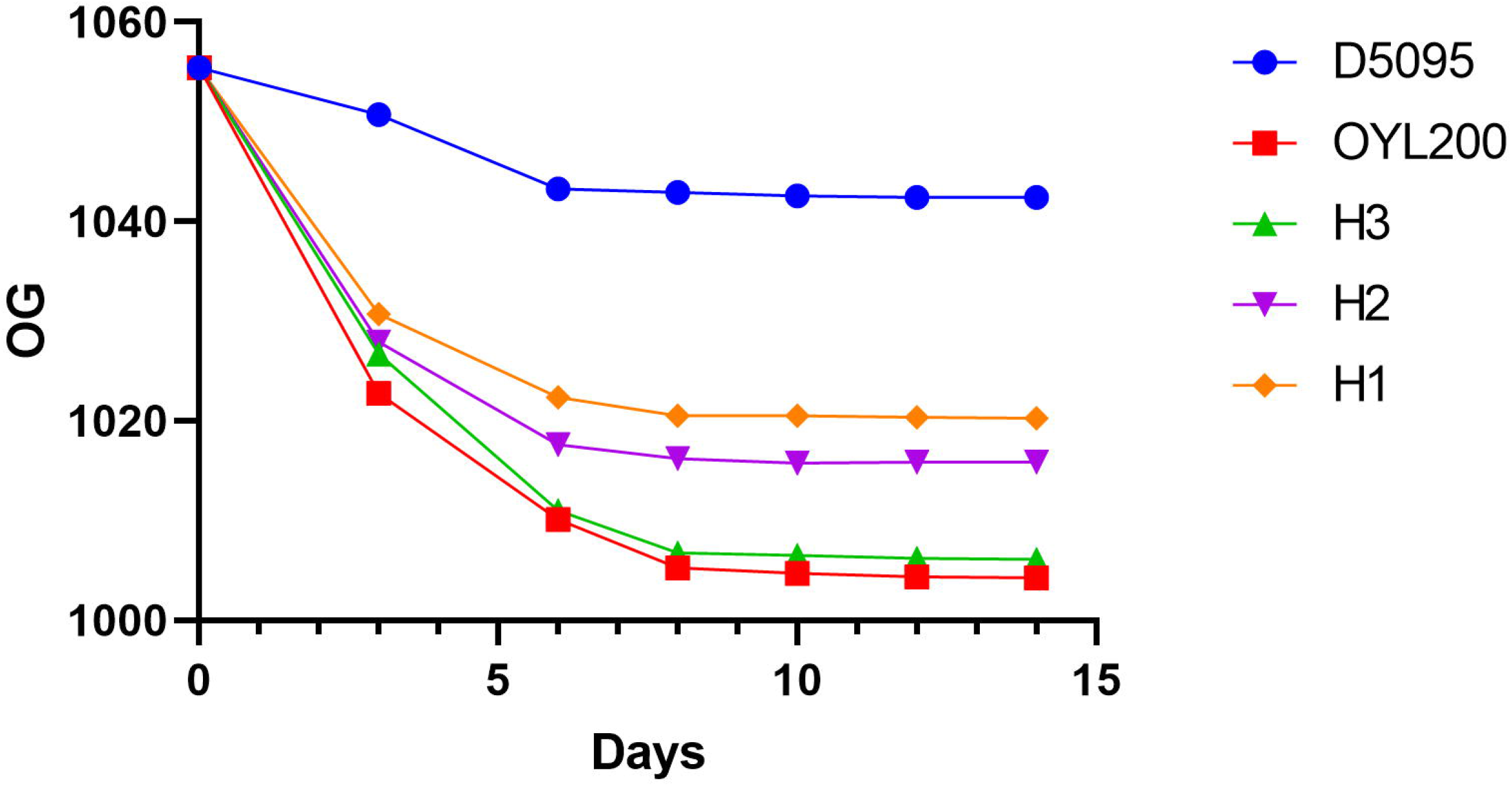
Fermentation kinetics of strains Sj-95, Sc-200 and the generated hybrids H1-3. Beer fermentation was done with pale wort of initial OG 1055.4. The plot contains reduction of sugars as OG (original gravity points) measured with density meter.

End point fermentation samples were analysed via HPLC to estimate the ethanol produced by the different strains (Fig. 4). The consumed sugars detected at the end of the fermentation confirmed the attenuation observed in the gravity data (Fig. 3). *S. jurei* D5095 showed the highest amount of residual sugars and the lowest production of ethanol compared to the Sc-200 and the generated hybrids. The ethanol produced (% ABV) varied between all strains, 6.6% (Sc-200), 6.4% (H3), 5.2% (H2), 4.5% (H1) and 1.8% (Sj-95).

**Figure 4:**
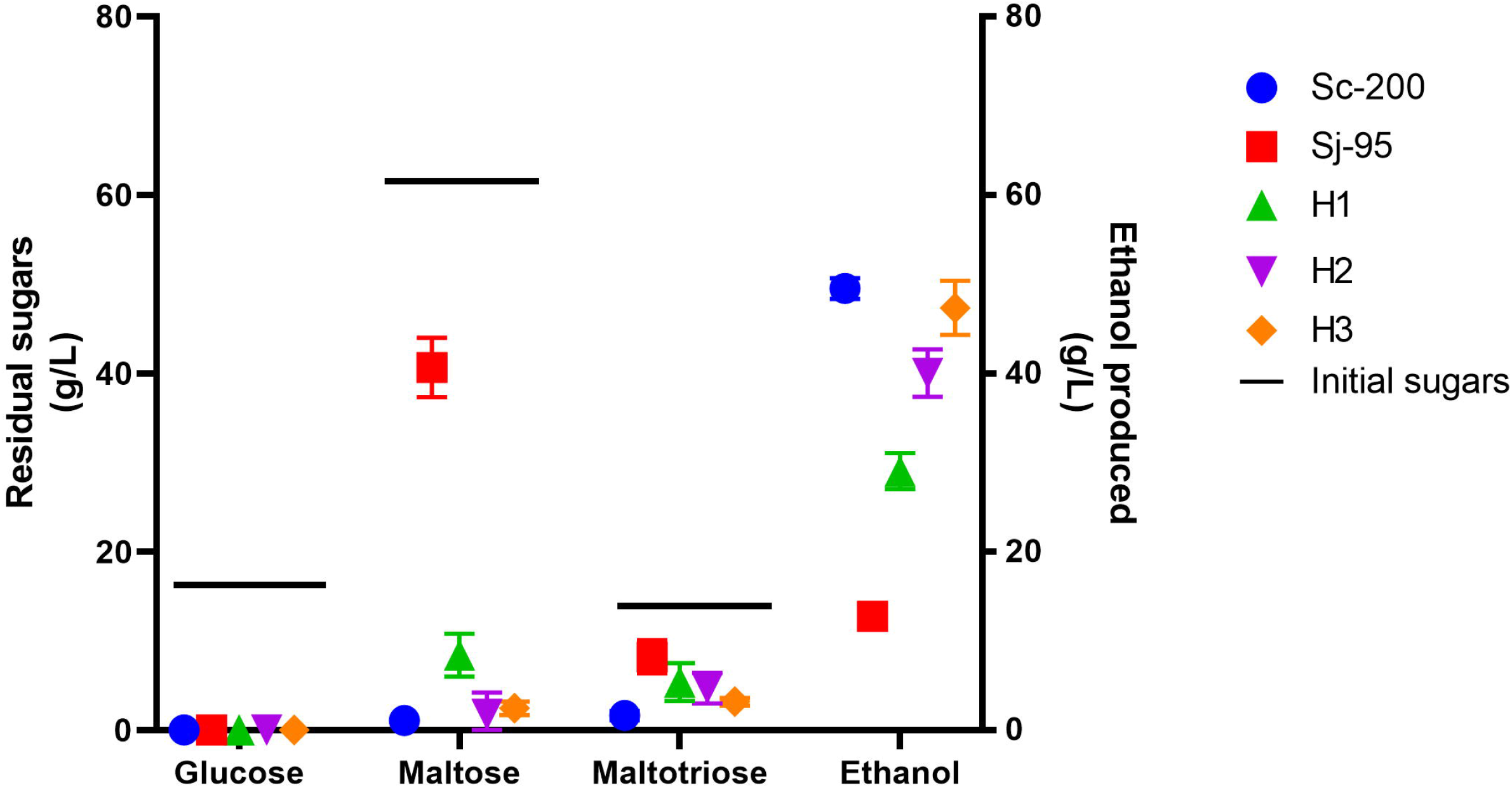
Sugar content and production of ethanol in final beer samples analysed via HPLC. Graph shows the residual wort sugars and the ethanol production at the end of fermentation from strains Sc-200, Sj-95, H1, H2, and H3.

Based on the hyper-attenuation characteristic that was observed in the fermentation profiles, we also tested for the presence of the *STA1* gene in the newly made hybrids via PCR. The results showed that the ale parent strain Sc-200 and H3 possess the *STA1* gene, while H1 and H2 do not (Fig. S4). It is likely that Sc-200 is heterozygote for the *STA1* gene, and H3 is the only hybrid that inherit that allele. In H1 and H2 the undesirable trait for hyper-attenuation has been eliminated.

### 3.4. Aroma profiling of generated hybrids reveals different aroma compounds from both S. jurei and S. cerevisiae parents

Identification of volatile aromatic compounds for all strains was carried out at the end of the beer fermentation using GCxGC-MS. In total, 18 esters, 11 alcohols and 33 volatile compounds including acids, terpenes, aldehydes, ketones and phenolic compounds were identified (Table 1). Peak area was used as an indication of relative compound concentration between beer samples.

**Table 1:** Target volatile metabolites of beer samples after 14 days of fermentation with Sj-95, Sc-200, H1, H2 and H3.

| **Esters** |  |  |  |  |  |  |
| --- | --- | --- | --- | --- | --- | --- |
|  |  | Peak Area x 10<sup>6</sup> |  |  |  |  |
| Compound name | Flavour description | Sj-95 | H1 | H2 | H3 | Sc-200 |
| Ethyl Acetate | fruity,pear | 0.10 | 14.07 | n.d | 2.17 | 20.55 |
| Isoamyl acetate | fruity,banana | 113.09 | 118.07 | 113.51 | 33.91 | 195.70 |
| Isobutyl acetate | fruity | 1.34 | 3.08 | 0.80 | 1.24 | 2.47 |
| Phenylethyl Acetate | floral,roses | n.d | n.d | 49.89 | 73.73 | 204.76 |
| Ethyl hexanoate | fruity,apple | n.d | 139.98 | 0.63 | n.d | 182.91 |
| Ethyl benzoate | medicinal | 0.10 | n.d | 0.83 | 0.13 | 1.31 |
| Propyl acetate | fruity,pear | 9.16 | 30.06 | 17.27 | 46.10 | 13.12 |
| Ethyl Propionate | fruity, pineapple | 5.91 | 12.57 | 5.93 | 16.99 | 5.79 |
| Ethyl heptanoate | fruity,grape | 9.52 | 14.08 | 23.08 | 8.11 | 16.54 |
| Ethyl octanoate | fruity,wine, sweet | 122.15 | 384.92 | 341.67 | 91.09 | 256.04 |
| Isopentyl isobutyrate | fruity,apricot, banana | 7.12 | 10.87 | 8.37 | 7.30 | n.d |
| Ethyl decanoate | sweet,apple, grape | n.d | 135.00 | 20.73 | 2.73 | 9.26 |
| Ethyl isobutyrate | fruity | 0.65 | 0.82 | 0.55 | 1.46 | 0.79 |
| Methyl Phenylacetate | honey | 3.28 | 3.75 | 3.31 | 3.60 | 3.13 |
| cis-Carvyl acetate | minty | n.d | n.d | 0.91 | 0.75 | 0.75 |
| Propyl octanoate | coconut, cacao | 0.12 | 5.47 | 0.50 | n.d | 0.10 |
| **Alcohols** |  |  |  |  |  |  |
|  |  | Peak Area x 10<sup>6</sup> |  |  |  |  |
| Compound name | Flavour description | Sj-95 | H1 | H2 | H3 | Sc-200 |
| Isoamyl alcohol | fusel, alcoholic, fruity | 775.86 | 523.41 | 615.71 | 0.58 | 1.67 |
| Phenylethyl alcohol | floral,rose, honey | n.d | 141.54 | 102.47 | n.d | 199.83 |
| Butanol | solvent,fusel | 4.22 | 4.79 | 4.55 | 5.62 | 17.69 |
| Propanol | solvent,fusel | 58.09 | 113.88 | 0.27 | 84.11 | 40.09 |
| Isobutanol | etheral,wine | 31.05 | 35.44 | 18.22 | 27.29 | 31.40 |
| Furfuryl alcohol | sweet | n.d | 3.13 | 1.44 | 2.83 | 1.67 |
| Hexanol | pungent,fusel, fruity | 19.07 | 23.32 | 17.57 | 22.85 | 19.38 |
| Methionol | meaty,onion | 0.59 | 0.63 | n.d | 0.81 | 0.98 |
| **Acids** |  |  |  |  |  |  |
|  |  | Peak Area x 10<sup>6</sup> |  |  |  |  |

| Compound name | Flavour description | Sj-95 | H1 | H2 | H3 | Sc-200 |
| --- | --- | --- | --- | --- | --- | --- |
| Hexanoic Acid | sour,cheesy | 114.68 | 191.37 | 110.19 | 171.14 | 126.50 |
| Butanoic Acid | acidic, unpleasant | 11.08 | 3.88 | 7.04 | 5.46 | 5.81 |
| Isovaleric Acid | cheesy | 12.92 | 10.76 | 9.12 | 17.20 | 9.96 |
| Isohexanoic acid | fruity | n.d | 0.08 | 0.03 | 0.69 | 0.42 |
| Propanoic acid | pungent, acidic | 2.07 | 5.02 | 3.94 | 8.40 | 1.64 |
| Acetic Acid | sour, acidic | 32.58 | 0.68 | 19.70 | n.d | 0.68 |
| Octanoic Acid | rancid, cheesy | n.d | 315.80 | 173.67 | 186.57 | 116.58 |
| **Terpenes and other volatile compounds** |  |  |  |  |  |  |
|  |  | Peak Area x 10<sup>6</sup> |  |  |  |  |
| Compound name | Flavour description | Sj-95 | H1 | H2 | H3 | Sc-200 |
| 1-alpha-Terpinyl acetate | herbal | 3.70 | 4.44 | 2.16 | 4.61 | 2.91 |
| Hop ether | floral,woody | 11.40 | 10.06 | 10.93 | 14.16 | 8.13 |
| Myrcene | woody,spicy | 3.79 | 14.06 | 13.58 | 9.78 | 10.76 |
| Linalool | citrus,floral, waxy | 97.57 | n.d | n.d | 107.32 | n.d |
| 4-vinyl guaiacol | clove,spicy, smoke | 1.95 | 2.30 | 1.36 | n.d | n.d |
| 2-4-di-tert-butylphenol | phenolic | 3.16 | 3.84 | 2.94 | 3.52 | 3.46 |
| Alpha-Phellandrene | citrusy, peppery | 0.56 | 0.63 | 0.58 | 0.58 | 0.68 |
| o-Cymene | spicy | 9.48 | 9.30 | 8.54 | 10.72 | 7.83 |
| Dehydro-p-cymene | fresh, citrus | 0.46 | 0.56 | 0.32 | n.d | 0.39 |
| Nonanol | citrus | 17.61 | 14.38 | 48.42 | 23.02 | n.d |
| Limonene | citrus, fresh, sweet | 0.61 | 0.16 | 0.90 | n.d | 0.27 |
| 2-Acetylfuran | sweet, nutty | n.d | 1.31 | n.d | 1.54 | 1.24 |
| 2-Hexanone | acetone like | 0.03 | 0.06 | 0.04 | n.d | 0.06 |
| Terpinolene | pine | 0.27 | n.d | 0.29 | 0.41 | 0.42 |
| Camphene | fresh,herbal,woody | 0.04 | 0.04 | 0.02 | 0.05 | n.d |
| (E)-2,6-Dimethylocta-1,5,7-trien-3-ol | camphoreous,lime | 1.73 | 2.29 | 1.17 | 2.01 | 2.44 |
| Nonanal | citrus | 0.31 | 0.54 | 0.55 | n.d | 0.54 |
| Ocimene | fruity | 13.09 | 11.71 | 9.43 | n.d | 9.77 |
| Pinene | woody,pine | n.d | 2.16 | n.d | 17.93 | 2.16 |
| 1-Decene | sweet | 0.29 | 5.06 | 7.55 | n.d | 5.79 |

As expected, the levels of volatile compounds varied between the parent strains. A strong tropical and fruity character was detected in Sc-200 due to the greater production of esters and other volatiles (Table 1). Sj-95 had instead a lower production of esters, but produced a significant amount of spice/clove and alcoholic aromas deriving from phenolic compounds and fusel alcohols, respectively. The hybrid strains inherited different complementary flavour profiles somewhere in between the parental strains. Ethyl hexanoate, a compound with a sweet fruity/apple aroma, derived from strain Sc-200 was only inherited in H1 and H2 hybrids. Compared to the parents, a higher production of specific compounds, such as ethyl propionate and α-terpinyl-acetate, which intensify some aroma characteristics, were also present in the hybrid. 4-vinyl-guaiacol, a spicy phenolic aroma was detected in Sj-95 and inherited in lower concentrations in H1 and H2, while not detectable in H3. A low production of propyl acetate and ethyl isobutyrate, both compounds with fruity aroma, was detected in both parental strains. Similar moderate production of the same metabolite is recorded in hybrids H1 and H2 but the beer made by H3 did show a higher concentration of this specific compound outperforming both parental strains. Different parental allele inheritance in the hybrids is responsible for the diversification of the aroma characteristics in the beer by strengthening or weakening specific features. Clustering of the samples according to the amount of compounds present in the fermentation products revealed the phenotypic relationships between the strains (Fig. 5). H1 and H2 aroma profile resembled those of Sc-200 and Sj-95, respectively. Interestingly, H3 does not clearly cluster with any of the two parents and show a more diverse flavour profile.

**Figure 5:**
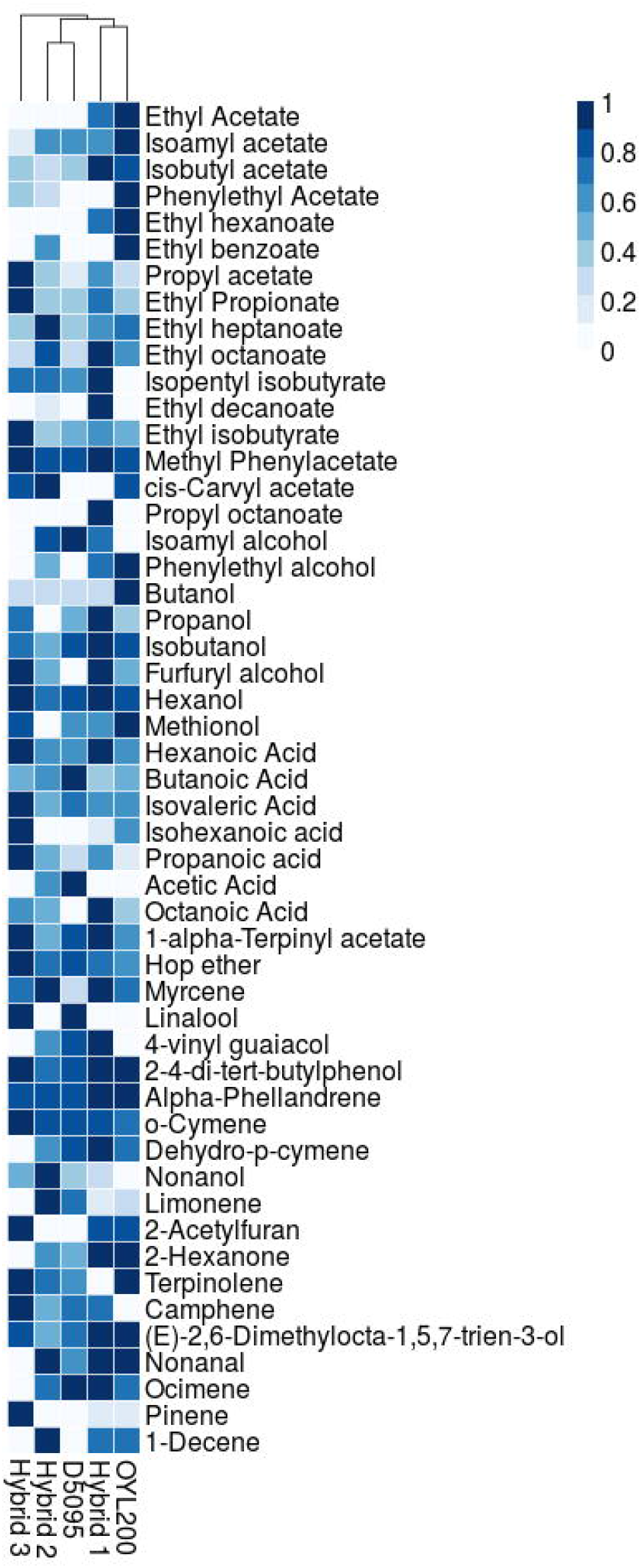
Clustering and heatmap visualization of aroma profiling. Normalized values per detected volatile compound across 14 days fermentation beer samples with Sj-95, Sc-200, H1, H2 and H3.

A sensory evaluation of the final beers was carried out at Cloudwater Brew Co (Fig. 6; Table S2). Finished beers made with H1 and H2 possess a nice estery character with predominant hits of apple, banana and pear. Metabolites related with these aromas were in fact detected in high concentrations through GC – MS in those beer samples (Table 1). A clove/phenolic character present in different intensities was tasted in both H1 and H2 beers and can be linked to the presence in low level of 4-vinyl-guaiacol. Beer made from H3 was described as fruity and tropical, with notes of peach and berries, with a more astringent and light sour character than H1 and H2, and with very low phenolic content. H2 aroma and flavour profile was considered as the most balanced one, with a nice sweetness and soft body characteristics. H1 and H2 were selected for larger scale beer fermentation in 200L barrels to assess the character in a more complex environment with the addition of hops.

**Figure 6:**
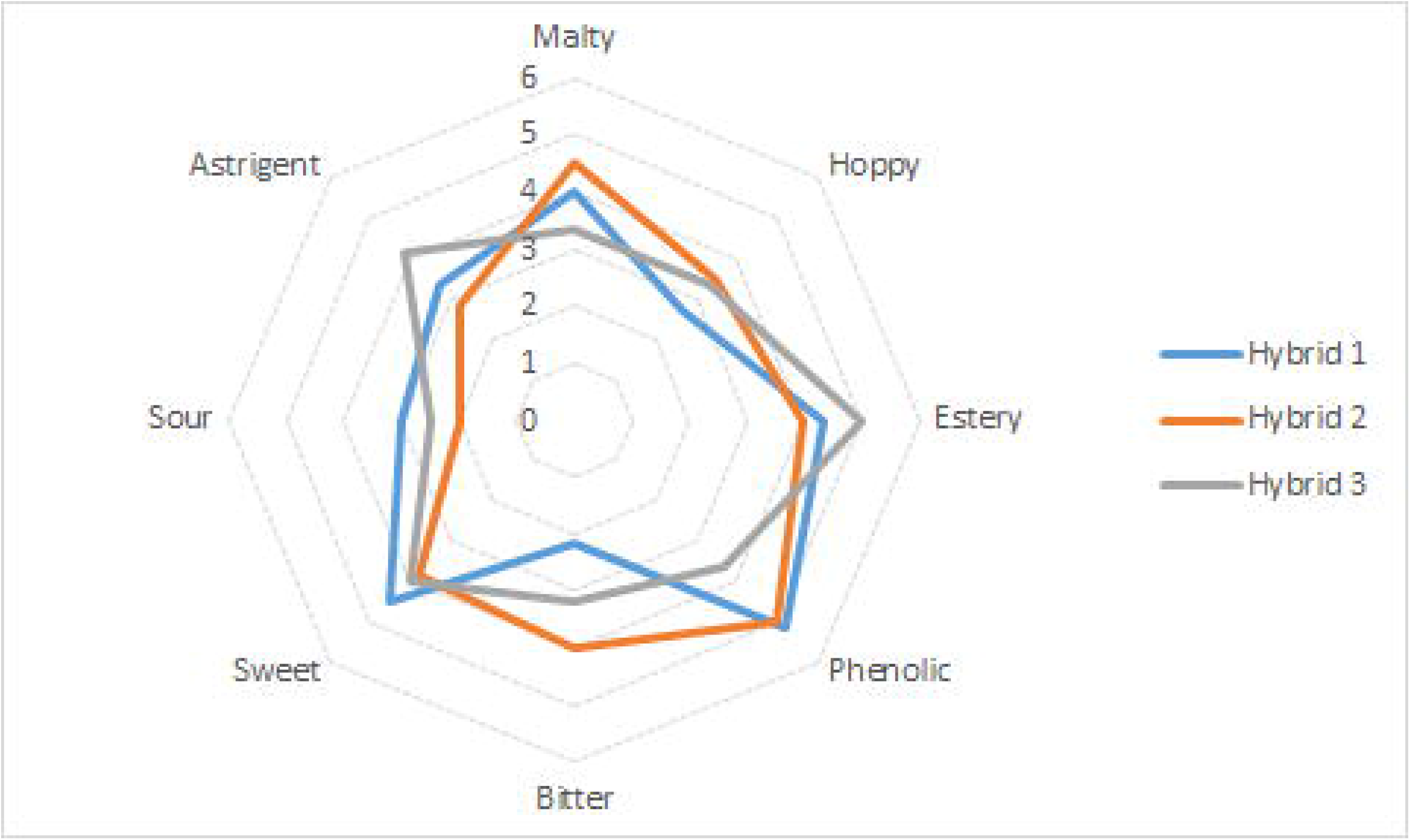
Descriptive sensory analysis. Final samples from 10L beer fermentation with Hybrids 1-3 for flavour assesment.

## 4. Conclusions

Recent years have brought a remarkable increase on the demand of craft beer. Consumers request unique recipes and flavours beyond the traditional and well known beer styles. Aromatic diversity and combination of different flavours is a way to create new products. Through interspecific hybridisation, complex aromatic profiles can be generated from the different parental species.

In this study, we evaluated the beer fermentation performance and aroma profile of *S. jurei* and non GMO generated brewing hybrids. *S. jurei* is able to utilize maltose and it can grow in low temperatures and this ability gives the potential of industrial applications and breeding with other brewing strains. We confirmed that *S. jurei* can also assimilate maltotriose, another brewing related sugar. It was shown that, *S. jurei*fermentation ability is limited when compared with a *S. cerevisiae* ale strain. Then, we constructed hybrids between those 2 different species through spore to spore mating. In that way, we can see a combination of inherited traits in the hybrids contributing to different phenotypes and growth characteristics. This is shown for several industrial relevant traits such as growth in maltose and maltotriose, which contribute to the good performance in brewing wort, and in growth at cold temperature. Also, the superior fermentation performance is observed in generated hybrids, overcoming *S. jurei* inability to complete the fermentation. This suggests that efficient wort sugar utilization is not essential to be deriving from both parental strains to create successful hybrids for the brewing industry and opens the potential on usage of alternative species in the production of fermented beverages.

Different allele inheritance in the generated hybrids resulted also in the elimination of the unwanted hyper-attenuation character. The *S. cerevisiae* ale parent is suspected to be heterogyzous to the particular gene conferring this characteristic. Therefore, through spore to spore mating the character is not present in all the hybrids. This is the first time that the diastatic activity of some *S. cerevisiae* strains can be eliminated through breeding techniques.

Also, in this study we performed aroma analysis on fermented beers from a selection of strains generated in this project. This is the first time the aroma profile of *S. jurei* in beer products is described. A combination of tropical and floral character was inherited in the hybrid strains deriving from complementary aroma profiles of both *S. jurei* D5095 and *S. cerevisiae* OYL200. The 3 hybrids present different concentrations and intensities of various compounds and complete absence or presence of some volatiles related with their parents. Loss of 4-vinyl guaiacol production is a signature of domestication in brewing yeast (Gonçalves et al., 2016) (Gallone et al., 2016) and it is indicated that the POF character can be eliminated through spore clone selection. Mixed or intermediary flavour profiles are achievable and highly recommended to facilitate yeast selection in specific beer styles or new recipes.

Applying classical breeding to ale strains can be challenging as strain often suffer from poor sporulation ability and efficiency (Codon et al., 1995) but interspecific hybridisation can be a way of creating new yeast strains conferring characteristics from different species. Various studies demonstrated hybridisation as a mechanism of domestication in beer and wine environments (Gallone et al., 2018) (Gallone et al., 2019)(Giannakou et al., 2020) (González et al., 2008) and also hybrids within the alternative *Saccharomyces* genus have been reported for their usage in fermented beverages (Krogerus et al., 2018) (García-Ríos et al., 2019) (Mertens et al., 2015). Selection of parental strains for breeding or hybridisation experiments possessing desirable characteristics will improve the phenotypic diversity of industrial candidates.

## Author contributions

DD conceived the study; DD and UV supervised the genetic and phenotypic experiments, and the GC-GC-MS analysis, respectively. MC and PJ supervised the pilot-scale fermentation. KG carried out all the experimental work with the inputs of FV. NN contributed to the construction of five hybrids. KG and FV analysed the genetic and physiological data and KG, FV and PZ performed and analysed the GC-GC-MS volatile compounds spectra. KG, FV and DD wrote the manuscript with input of MC, PZ, UV. All authors contributed to the article and approved the submitted version.

## Acknowledgments

We thank Silvia Carlin for the assistance during the GC × GC-MS analysis and Daniel Schindler for the Omega yeast collection. We thank the production team of Cloudwater Brew co for helpful advice on the brew making and diastaticus yeast.

## Supplementary Table Legends

**Table S1**: Specific growth rate (μmax) of strains D5095, D5088, OYL200, OYL500 and H1-8 in YP-maltose, YP-maltotriose, YPD and DME for 72h.

**Table S2**: Overview results sensory evaluation from 10L beer fermentation by professional tasting panel. Score legend: absent (0); very very low presence (1); very low presence (2); low presence (3); mild presence (4); presence (5); clear presence (6); strong presence (7)

## Supplementary Figure Legends

**Figure S1**: **Growth kinetics of *S. jurei* D5095, D5088, *S. cerevisiae* ale strain OYL200, type strain NCYC505 and natural isolate 96.2.** Panel A describes the growth in maltose; panel B describes the growth in maltotriose. Plate reader cultivations with online monitored optical density for 54h.

**Figure S2**: **Species-specific PCR confirmation of successful hybrid construction in 1.5% agarose gel.** Panel A: Lane 1 100 bp ladder, lane 2 D5095, lane 3 OYL200, lanes 4-6 Hybrids H1-3 containing DNA from both parents Panel B: M: 100bp ladder, Lanes 1 and 3: D5088 and OYL500, lanes 5,7,8,10 and 11 contain genomic DNA from H4, H5, H6, H7 and H8 respectively.

**Figure S3**: **Fluorescence flow cytometry analysis**. Top row left to right; Control ploidy strains D5095 (diploid), BY4742 (haploid) and PB7 (tetraploid). Bottom row left to right; Hybrid strains H1, H2 and H3.

**Figure S4**: **Detection of *STA1* gene via PCR**. Lanes 1-4: strains Sc-200, H1, H2, and H3, respectively. M: marker, Hyperladder 1kb.

