## Supplementary materials for "Biotechnological exploitation of *Saccharomyces jurei* and its hybrids in craft beer fermentation uncovers new aroma combinations"

Supplementary material for *S. jurei* paper

**Table S1**: Specific growth rate (μmax) of strains D5095, OYL200, H1, H2 and H3 in DME for 72h.

| | | Sj-95 | Sc-200 | H1 | H2 | H3 | Sj-88 | Sc-500 | H4 | H5 | H6 | H7 | H8 |
|---|---|---|---|---|---|---|---|---|---|---|---|---|---|
| YPD | Specific growth | 0.06 ± 0 | 0.07 ± 0.01 | 0.06 ± 0 | 0.07 ± 0 | 0.07 ± 0 | 0.061 ± 0.013 | 0.056 ± 0.004 | 0.047 ± 0.018 | 0.078 ± 0.005 | 0.041 ± 0.007 | 0.075 ± 0.016 | 0.066 ± 0.016 |
| | Lag phase | 7.49 ± 0.85 | 9.57 ± 1.31 | 8.81 ± 1.21 | 9.89 ± 0.94 | 10.4 ± 0.4 | 9.52 ± 1.01 | 8.8 ± 2.39 | 10.45 ± 2.81 | 23.88 ± 2.8 | 16.48 ± 3.71 | 19.22 ± 3.2 | 14.58 ± 5.99 |
| | Maximum biomass | 1.32 ± 0.01 | 1.43 ± 0.01 | 1.37 ± 0.01 | 1.36 ± 0 | 1.44 ± 0.01 | 1.16 ± 0.02 | 1.18 ± 0.04 | 0.93 ± 0.04 | 1.04 ± 0.05 | 0.79 ± 0.07 | 1.09 ± 0.08 | 1.12 ± 0.08 |
| | Integral area | 74.1 ± 0.76 | 78.47 ± 0.22 | 74.39 ± 0.16 | 74.71 ± 0.92 | 79.58 ± 0.81 | 56.68 ± 2.93 | 58.87 ± 1.51 | 42.06 ± 4.52 | 44.13 ± 4.18 | 37.17 ± 3.05 | 49.42 ± 5.88 | 54.17 ± 5.48 |
| YP + 2% maltose | Specific growth | 0.032 ± 0.002 | 0.061 ± 0.002 | 0.065 ± 0.003 | 0.043 ± 0.003 | 0.047 ± 0.012 | 0.017 ± 0.003 | 0.025 ± 0.004 | 0.034 ± 0.001 | 0.023 ± 0.001 | 0.018 ± 0.001 | 0.016 ± 0.003 | 0.031 ± 0.001 |
| | Lag phase | 32.73 ± 1.59 | 15.09 ± 0.73 | 16.51 ± 0.74 | 20.87 ± 1.51 | 16.23 ± 1.11 | 2.12 ± 1.22 | 9.54 ± 1.65 | 12.88 ± 0.76 | 11.62 ± 1.93 | 19.94 ± 3.86 | 8.11 ± 4.47 | 12.84 ± 1.67 |
| | Maximum biomass | 1.2 ± 0.018 | 1.15 ± 0.004 | 1.17 ± 0.03 | 1.19 ± 0.041 | 1.24 ± 0.029 | 0.6 ± 0.02 | 1.05 ± 0.01 | 1.01 ± 0.01 | 1.08 ± 0.02 | 0.87 ± 0.05 | 0.88 ± 0.1 | 1.01 ± 0.02 |
| | Integral area | 31 ± 0.69 | 55.07 ± 0.85 | 54.68 ± 0.83 | 44.59 ± 0.18 | 49.72 ± 0.31 | 24.31 ± 1.92 | 38.96 ± 3.53 | 43.19 ± 0.81 | 37.33 ± 0.96 | 26.8 ± 1.99 | 29.08 ± 2.31 | 42.07 ± 1.19 |
| YP + 2% maltotriose | Specific growth | 0.02 ± 0.001 | 0.05 ± 0 | 0.03 ± 0.003 | 0.05 ± 0.003 | 0.03 ± 0.001 | 0.03 ± 0.003 | 0.035 ± 0.008 | 0.029 ± 0.004 | 0.02 ± 0.004 | 0.022 ± 0.008 | 0.018 ± 0.001 | 0.024 ± 0.005 |
| | Lag phase | 5.83 ± 1.38 | 13.66 ± 0.56 | 14.84 ± 1.17 | 14.19 ± 0.1 | 12.79 ± 1.85 | 3.06 ± 0.64 | 6.87 ± 1.39 | 8.13 ± 2.23 | 2.83 ± 1.51 | 7.83 ± 3.77 | 0.43 ± 0.29 | 2.32 ± 1.61 |
| | Maximum biomass | 0.68 ± 0.005 | 1.21 ± 0.015 | 1.2 ± 0.009 | 1.16 ± 0.006 | 1.29 ± 0.018 | 0.55 ± 0.04 | 1.16 ± 0.02 | 1.08 ± 0.05 | 0.6 ± 0.07 | 0.94 ± 0.17 | 0.66 ± 0.02 | 0.63 ± 0.08 |

| | | | | | | | | | | | | | |
|---|---|---|---|---|---|---|---|---|---|---|---|---|---|
| | Integral area | 24.74 ± 1.42 | 55.27 ± 0.69 | 45.96 ± 0.51 | 54.05 ± 0.22 | 51.19 ± 0.46 | 26.56 ± 1.99 | 49.97 ± 1.21 | 46.49 ± 2.6 | 22.15 ± 4.32 | 37.62 ± 10.69 | 22.32 ± 1.71 | 26.09 ± 3.49 |
| DME | Specific growth | 0.1309 ± 0.013 | 0.1475 ± 0.005 | 0.1265 ± 0.003 | 0.1385 ± 0.012 | 0.1139 ± 0.009 | 0.096 ± 0.01 | 0.099 ± 0.01 | 0.105 ± 0.02 | 0.109 ± 0.01 | 0.144 ± 0.04 | 0.107 ± 0.01 | 0.09 ± 0.01 |
| | Lag phase | 15.92 ± 15.97 | 6.79 ± 0.58 | 13.09 ± 8.13 | 8.48 ± 1.81 | 6.69 ± 0.31 | 5.23 ± 1.56 | 15.96 ± 7.47 | 19.59 ± 14.93 | 32.41 ± 15.19 | 9.1 ± 3.07 | 8.51 ± 2.82 | 4.79 ± 0.9 |
| | Maximum biomass | 85.21 ± 3.21 | 74.07 ± 3.27 | 65.76 ± 7.02 | 79.63 ± 4.27 | 80.27 ± 11.28 | 81.45 ± 10.33 | 134.61 ± 15.97 | 98.52 ± 9.12 | 103.91 ± 19.69 | 137.27 ± 44.21 | 120,41 ± 23.85 | 79.02 ± 6.97 |
| | Integral area | 3668 ± 91 | 3264 ± 127 | 2856 ± 152 | 3533 ± 147 | 3146 ± 355 | 3736 ± 456 | 5173 ± 599 | 4504 ± 629 | 3953 ± 643 | 5310 ± 1397 | 4746 ± 568 | 3412 ± 454 |

**Table S2**: Overview results sensory evaluation from 10L beer fermentation by professional tasting panel. Score legend: absent (0); very very low presence (1); very low presence (2); low presence (3); mild presence (4); presence (5); clear presence (6); strong presence (7)

| Beers made with strain: | | Hybrid 1 | Hybrid 2 | Hybrid 3 |
|---|---|---|---|---|
| **Aroma/Flavour** | Malt | 4 | 4.5 | 3.5 |
| | Hoppy | 3 | 3.5 | 3.5 |
| | Esters | 4.5 | 4 | 5 |
| | Phenolic | 5.5 | 5 | 4 |
| **Comments** | | Fruity, pear, grape, lemon, estery, sweet, pineapple, banana, slightly sour, vanilla, phenolic, clove, low sulphur | Ethyl acetate, acetaldehyde, isoamyl acetate, estery, pear, light hoppy, sweet, soft, nice body, phenolic, funky, clove, not very carbonated | Lemon, spicy, green apple, dry, low phenolic, light hoppy, light bitter, peach, red berry, yeasty, light sour |
| **Taste/mouthfeel** | Bitterness | 2.5 | 4 | 3.5 |
| | Sweet | 4.5 | 3.8 | 4 |
| | Body | 3 | 3 | 2 |
| | Astringency | 3.5 | 3 | 4 |
| | Sour | 3 | 2 | 2.5 |
| **Overall impression** | Appreciation | 6 | 6 | 5 |
| | Complexity | 4 | 5 | 5 |
| | Balance | 5 | 4.5 | 4 |

**Table S3**: Pale wort recipe

| **Malts** | % |
|---|---|
| Golden Promise | 74.8% |
| Flaked Oats | 13.7% |
| Wheat | 9.2% |
| Caramel pils | 2.3% |
| **Mashing profile** | Time |
| 50 °C | 15 min |
| 72 °C | 60 min |
| 77 °C | 5 min |
| 100 °C (boiling step) | 60 min |

**Figure S1**: **Growth kinetics of *S. jurei* D5095, D5088, *S. cerevisiae* ale strain OYL200, type strain NCYC505 and natural isolate 96.2.** Panel A describes the growth in maltose; panel B describes the growth in maltotriose. Plate reader cultivations with online monitored optical density for 54h.

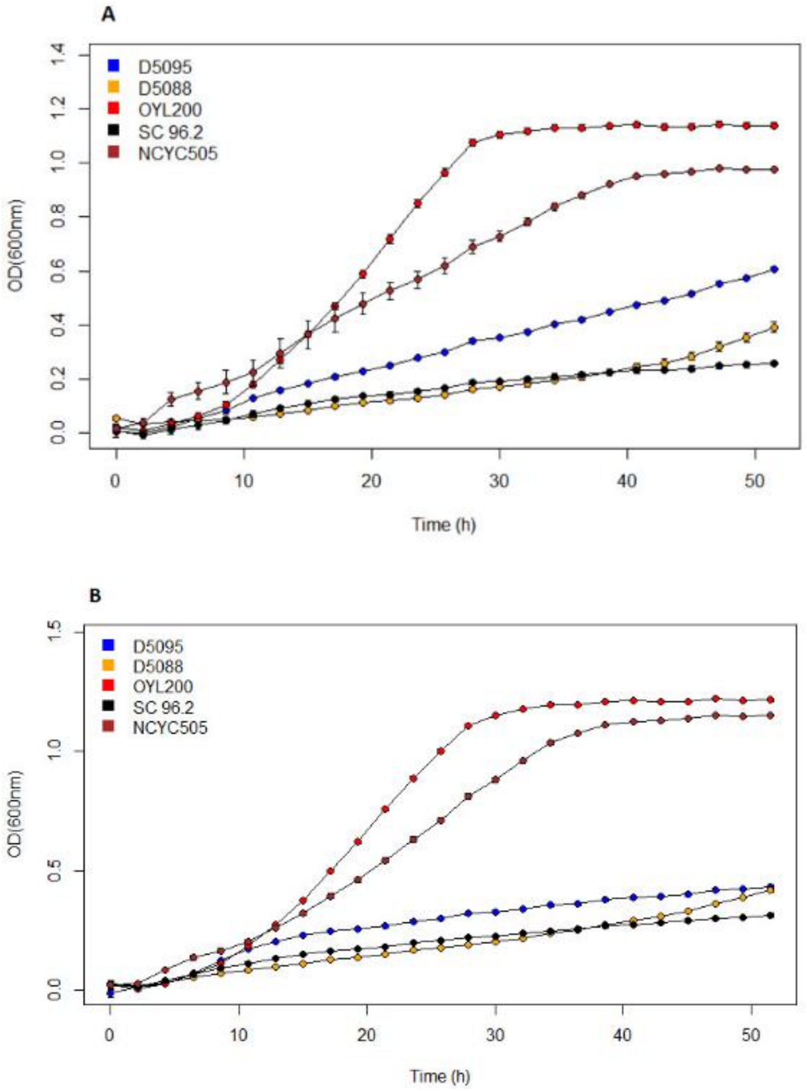

**Figure S2**: **Species-specific PCR confirmation of successful hybrid construction in 1.5% agarose gel.** Panel A: Lane 1 100 bp ladder, lane 2 D5095, lane 3 OYL200, lanes 4-6 Hybrids H1-3 containing DNA from both parents Panel B: M: 100bp ladder, Lanes 1 and 3: D5088 and OYL500, lanes 5,7,8,10 and 11 contain genomic DNA from H4, H5, H6, H7 and H8 respectively.

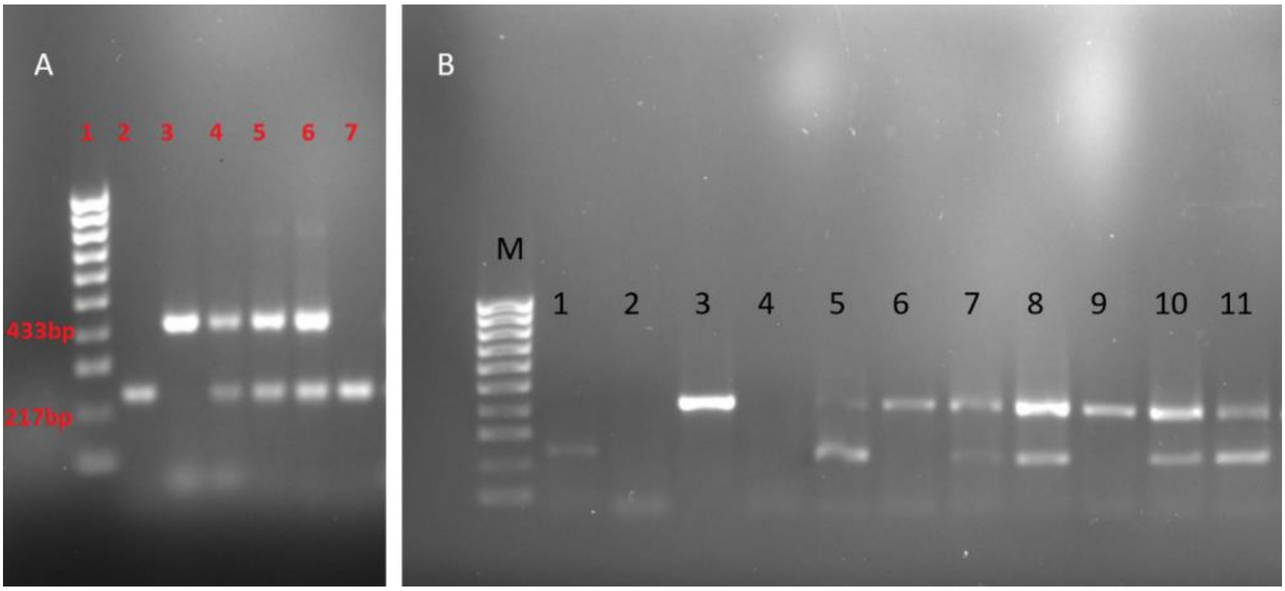

**Figure S3**: **Fluorescence flow cytometry analysis**. Top row left to right; Control ploidy strains D5095 (diploid), BY4742 (haploid) and PB7 (tetraploid). Bottom row left to right; Hybrid strains H1, H2 and H3.

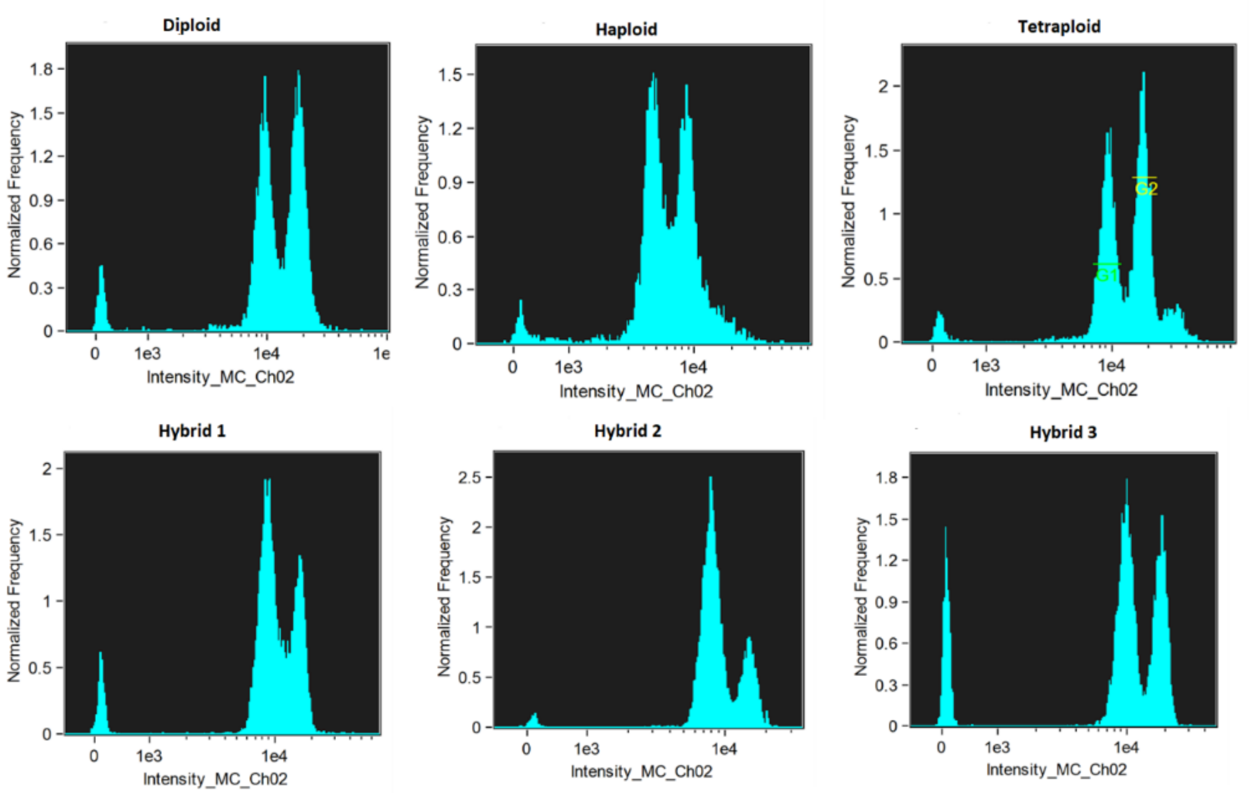

**Figure S4**: **Detection of *STA1* gene via PCR**. Lanes 1-4: strains Sc-200, H1, H2, and H3, respectively. M: marker, Hyperladder 1kb.

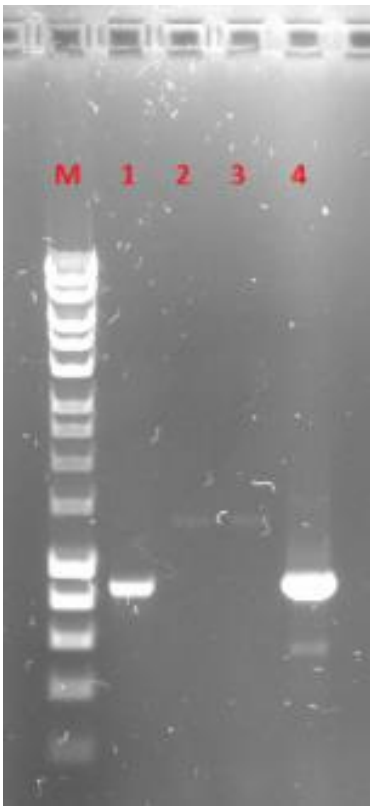
